# Microbial interactions within a community are insensitive to recurrent disturbances

**DOI:** 10.64898/2026.08.12.744439

**Authors:** M. Javad Vahramian, Renato Mirollo, Babak Momeni

## Abstract

Environmental disturbances are ubiquitous in microbial ecosystems, yet their effects on community composition and interactions remain poorly understood. Here, we combine analytical derivations and large-scale generalized Lotka–Volterra simulations to examine how recurrent disturbances, varying in intensity (*C*), frequency (1/*T*), regularity, and evenness, affect microbial communities. We find that context-dependent interactions (*I*_*CD*_)—representing the net effect of the entire community on each species—are robust to changes in disturbance intensity and frequency. Community composition is influenced by disturbance, but this effect is primarily captured by a single composite parameter *T*/log(*C*), reducing the two-dimensional disturbance space onto a one-dimensional axis. Species abundances and *I*_*CD*_ remain largely independent of specific values of *T* and *C* except near critical transition regions (survival thresholds and no-interaction zones). Sensitivity analyses show that disturbance-sensitive cases occur in the vicinity of these boundaries. This robustness extends to when the disturbance is irregular (even up to 50% stochastic variation in *T* or *C*), with uneven disturbances, when the disturbance is species-specific, and holds at any time point throughout the inter-disturbance growth phase. The patterns are independent of species richness, growth rates, carrying capacities, and interaction strengths. Our findings highlight that microbial community organization is largely conserved under recurrent environmental stress, with extinction acting as the dominant route by which disturbances reshape communities. These findings indicate that disturbance is not a strong driver of community structure and that minor variations in experimental or environmental conditions or sampling time are not expected to substantially impact microbiome composition.

## 1. Introduction

Environmental disturbances are pervasive in microbial ecosystems. Across natural and engineered habitats, microbial communities routinely experience fluctuations in temperature (Burman & Bengtsson-Palme, 2021; Fujikawa & Sakha, 2014), nutrient availability (Nguyen et al., 2021), hydration (Schimel, 2018), and other abiotic factors (Dedrick et al., 2021; Gallardo-Navarro et al., 2024). These disturbances vary widely in frequency and intensity (Berga et al., 2012; Shade et al., 2012), yet microbial communities often maintain relatively stable composition and function despite recurrent change (Philippot et al., 2021). This apparent robustness raises a question in microbial ecology: how do complex microbial communities preserve their structure and function in the face of recurrent disturbance?

Understanding how microbial communities respond to recurrent disturbances is important for both fundamental ecology and practical applications. Theoretically, it sheds light on the principles of stability and resilience in complex ecosystems (Philippot et al., 2021). Practically, it helps interpret experimental data, guides the design of microbiome studies, and improves predictions of microbial dynamics in variable natural environments where stochastic disturbances are common (Mancuso et al., 2021).

Despite substantial advances in microbial ecology, our understanding of community responses to recurrent disturbances remains limited, because of the scarcity of temporally resolved observational data (Buckley et al., 2021; Shade et al., 2012) and the challenges of experimentally controlling the disturbance regime (Banitz et al., 2020; Buma, 2021). Most observational studies provide either discrete snapshots or infrequent samples, offering little insight into dynamics between sampling points. In laboratory settings, it is technically challenging to control disturbance frequency, intensity, and regularity over extended periods. As a result, the dynamical behavior of microbial communities under recurrent environmental disturbances remains poorly characterized (Shade et al., 2012).

Mathematical and computational modeling offers a powerful way to overcome these empirical limitations (Song et al., 2014). Differential-equation-based frameworks, particularly generalized Lotka–Volterra (gLV) models (Stein et al., 2013; Momeni et al., 2017; Venturelli et al., 2018), provide a minimal yet analytically tractable description of microbial growth and interspecific interactions (Gonze et al., 2018). These approaches have delivered key insights into coexistence, colonization resistance (Kurkjian et al., 2021), invasion dynamics (Hu et al., 2025), and community assembly (Dal Bello et al., 2021; Lee et al., 2023), and they enable systematic *in silico* exploration of disturbance regimes that are difficult to manipulate experimentally.

In this study, we use a generalized Lotka-Volterra framework to systematically investigate how recurrent disturbances affect microbial community composition and within-community interactions. Our main result is that microbial communities exhibit striking robustness to variations in disturbance intensity and frequency. Changes in these parameters have minimal impact on stable community composition and interactions, except under conditions that approach species extinction thresholds. As a consequence, we conclude that extinction or proximity to extinction thresholds, rather than gradual shifts in interactions, is the primary mechanism by which recurrent disturbances reshape microbial communities.

## 2. Methods

### 2.1. Model Framework: Lotka-Volterra with Periodic Disturbance

The foundational framework for this study is the Lotka-Volterra (LV) model, which describes microbial population dynamics and interspecies interactions through a system of coupled differential equations. We use the following form of LV to study the dynamics of microbial communities:

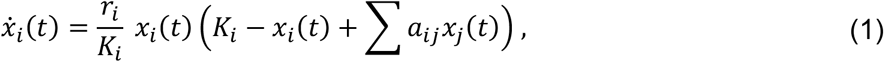

where *r*_*i*_, and *K*_*i*_ are the growth rate and carrying capacity of species *i*, and *a*_*ij*_ is the interaction strength of species *j* over species *i*. In its general form, the model assumes continuous population growth. To simulate recurrent environmental disturbances, we introduce periodic dilution events, which impose discontinuities in the system. These dilution events, occurring at regular or irregular intervals, reduce the microbial population by a factor of *C* > 1, where 1/*C* of the population remains after each disturbance. This key modification transforms the model from a continuous dynamical system into one punctuated by discrete disturbances, allowing for a more realistic representation of environmental fluctuations.

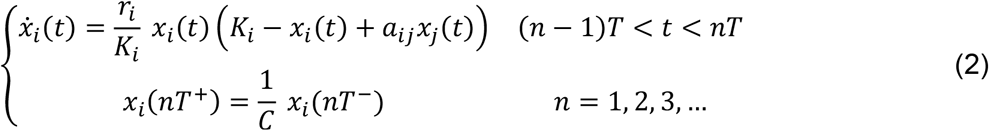

### 2.2. Model parameterization

The nasal microbiota was selected as the model microbial community for this study due to its relevance in low-nutrient environments, where the Lotka–Volterra model has previously been shown to perform well (Dedrick et al., 2021). Published data in our previous works related to the basal growth rates, carrying capacities, and pairwise interspecies interaction strengths of six species of nasal microbiota were used to determine the range of these parameters in simulated communities. For each microbial community we first pick the microbial species with random parameters: growth rate *r*_*i*_ = 0.3 ± 0.05*u*, carrying capacity *K*_*i*_ = 0.5 ± 0.2*u*, and interaction coefficients *a*_*ij*_ = −2 ± 2.4*u*, where *u* is a standard uniform random variable with a distribution *U*(0,1) and *i* and *j* indicate species indices. We start the community dynamics simulations at equal abundances for all species and let them grow under various disturbance conditions and record the composition after 100 generations, when the community is expected to have stabilized in composition.

### 2.3. Simulation Design and Analysis

For each simulation, a unique community was generated by picking species whose growth rates, interaction strengths, and carrying capacities were randomly drawn from the observed ranges of nasal bacterial species. While most simulations were performed with three-species communities, additional simulations involving different community sizes were conducted for comparison.

All simulations were initialized with all species at equal abundances. The system was simulated under a disturbance regime for 100 generations to ensure convergence to a stable equilibrium. Preliminary analyses indicate that most communities reach stability within 10 to 20 generations, but extended simulation time was used to ensure robustness across all scenarios. All simulations were implemented in Matlab (see Code Availability for access).

### 2.4. Theoretical Analysis of the Impact of Environmental Disturbance

The model consists of two distinct components: a continuous growth phase and a discrete dilution event. Incorporating these two processes requires slightly different analytical approaches. Accordingly, we organize our calculations into two parts, first addressing the continuous component, and then the discrete one. In both analyses, we proceed step by step. We begin with the simplest case of a single species and then extend the framework to a community of multiple species.

#### 2.4.1. Growth Phase of an Isolated Microbe

Under the assumptions of the Lotka–Volterra (LV) system, the dynamics during the growth phase depend solely on the initial population. We first analyze how the population growth depends on the initial conditions. The solution to this system is determined by the initial values *x*_0_. We denote the corresponding trajectory by 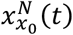, representing the population size of the species at time *t* during the *N*^*th*^ growth cycle. Notice that after each dilution event: 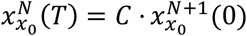 .

For a community consisting of a single microbial species isolated from any interactions with other species, we assume the following logistic-type equation:

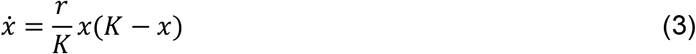

where r denotes the intrinsic growth rate and K the carrying capacity. Separating variables gives:

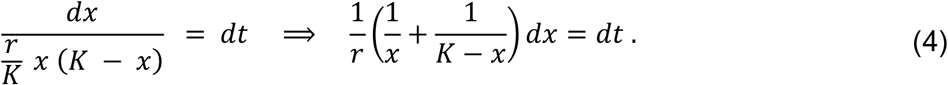

Integrating both sides yields

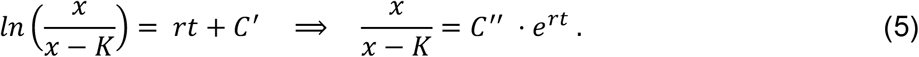

We can calculate the value of *C*^”^ by looking at the equations at *t* = 0:

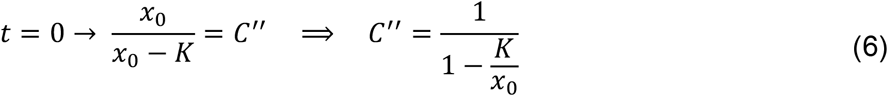

Now, solving for *x*(*t*) and assuming *x*_0_ ≠ 0 we obtain:

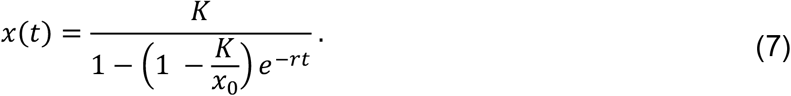

In the trivial case *x*_0_ = 0, the trivial solution *x*(*t*) = 0 is the only solution.

#### 2.4.2. Steady-State Abundance of an Isolated Microbe

To incorporate the effect of discrete dilution events into the continuous growth model, we treat each dilution as an instantaneous reduction in population size following a period of continuous growth. Based on Equation (7), the abundance starting at *x*_0_ and after the dilution step is

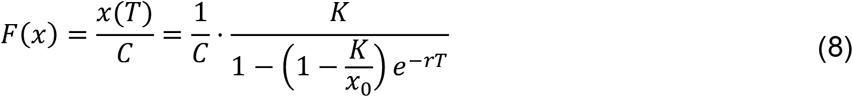

Setting *F*(*x*) = *x* gives us the equilibrium point 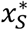:

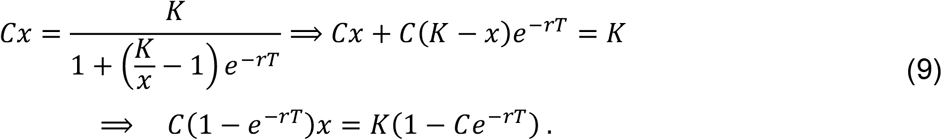

Thus, the steady state abundance, 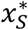, is equal to:

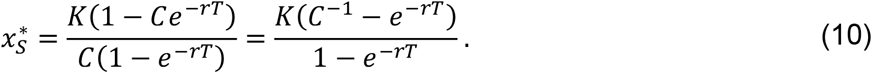

For the equilibrium composition, 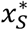, to be positive, we need to have *C*^−1^ − *e*^−*rT*^ > 0, or equivalently *Ce*^−*rT*^ < 1. Thus, for a community of a single species, if *Ce*^−*rT*^ ≥ 1, the only equilibrium state is 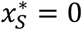 which is stable. Otherwise, if *Ce*^−*rT*^ < 1, there exists a non-trivial solution shown in equation (10).

In cases where a steady state solution exists, we can check its local stability by calculating the first derivative of *F*.

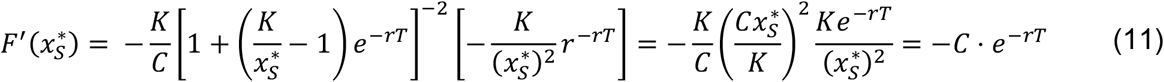

Equation (11) shows that 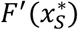 is strictly negative, indicating that the equilibrium is stable.

#### 2.4.3. Growth phase of a community of multiple species

We now consider a community of *n* weakly interacting species. The population dynamics in this case are governed by:

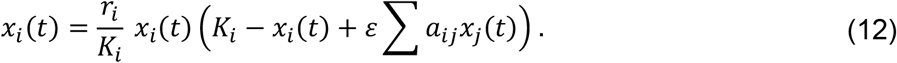

Notice that we are using the parameter *ε* to emphasize that for a small enough value of *ε*, the interaction coefficients can be made small compared to other parameters.

Using perturbation theory, we assume a solution of the form 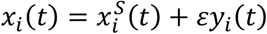, where 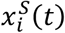 is the single-species solution. Substituting into the dynamic equation gives:

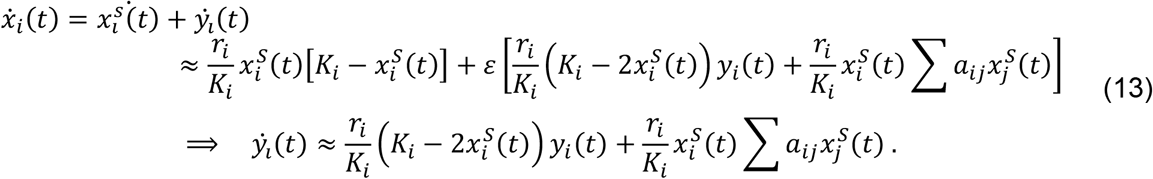

To solve for 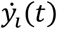, we introduce the integration factor 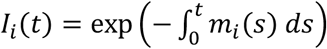 using 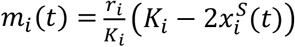 and 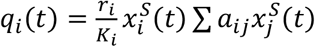. Thus,

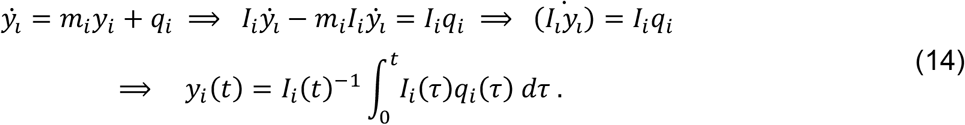

To calculate *I*_*i*_(*t*), we first expand *m*_*i*_(*t*) using equation (7) for 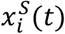, then integrate the resulting formula:

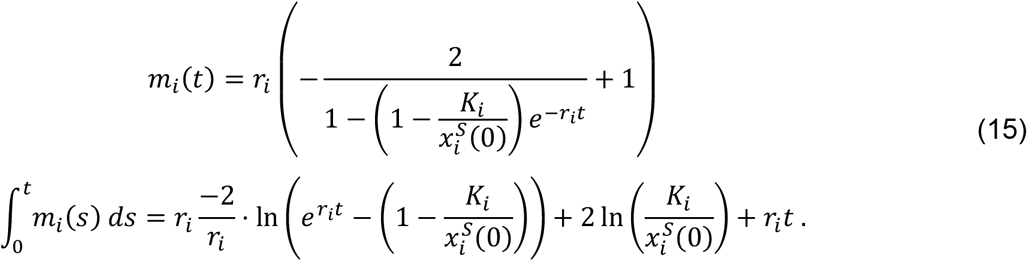

Similarly, for *I*_*i*_(*t*) we have:

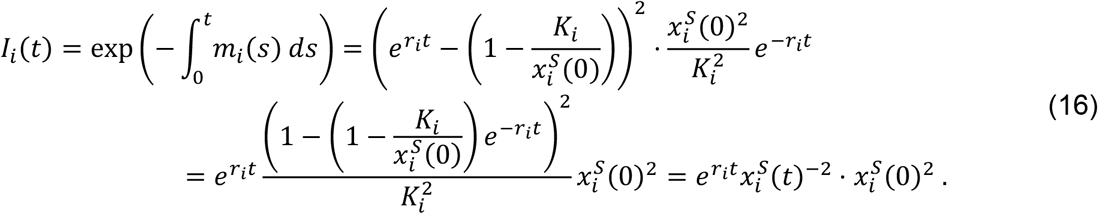

Expanding the last line of equation (14) using equation (16) gives the formula for *y*_*i*_(*t*) as:

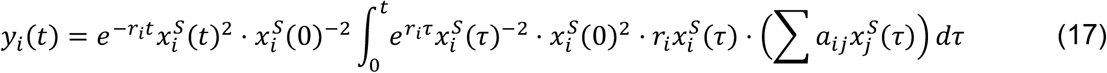

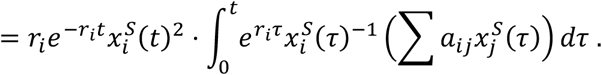

The resulting *y*_*i*_(*t*) represents changes in the population dynamics of species *i* because of interactions with others in the context of the community.

#### 2.4.4. Steady-State Abundance of a Microbial Community

To find the equilibrium populations, let *ε* > 0 and assume 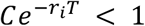 for all *i*. We write 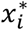, the steady-state population as a combination of two components:

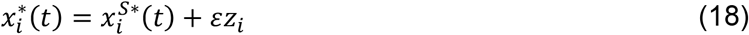

The first component represents the uncoupled steady-state (i.e., in the absence of any community interactions), and the second part is the first order perturbation approximation because of the impact of other community members. Let *X* = (*x*_1_, . . ., *x*_*n*_) and *Z* = (*z*_1_, . . ., *z*_*n*_) be the vector form of the stable states.

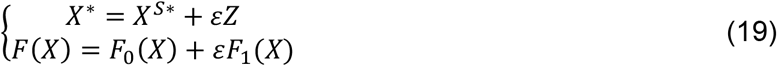

We aim to find the fixed point. This is equivalent to finding the solution to *F*(*X*^∗^) = *X*^∗^. Equations below hold up to the first order estimate around *X*^∗^.

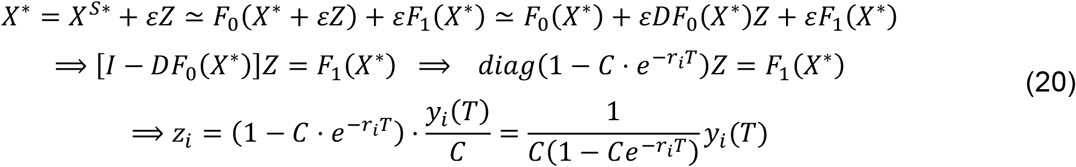

with the value of 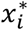 calculated in (10) and (17), respectively. Thus, in first-order approximation, the steady-state solution in a multi-species community consists of two components: one determined by the intrinsic single-species dynamics and another arising from interspecies interactions:

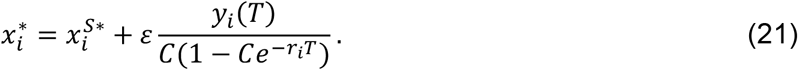

#### 2.4.5. The Case of Species with Similar Growth Rates

As part of our theoretical and perturbation analysis, we examine the special case of species that either grow identically or possess only small differences in their intrinsic growth rates. To formalize this, we represent the growth rate of species *r*_*i*_ = *r* + *ε*_*i*_, where *ε*_*i*_ is a Gaussian random variable of small magnitude relative to r. This formulation captures natural variability among species while preserving analytical tractability. In the case of identical growth rates, where all *r*_*i*_ = *r*, the intrinsic single-species component is proportional to carrying capacities, given by:

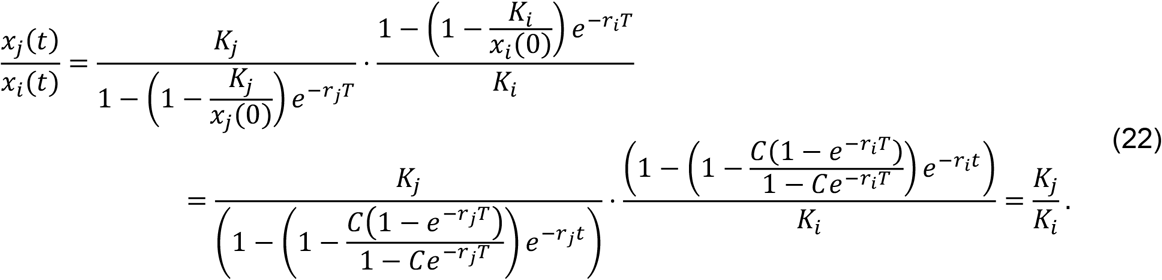

The interaction-driven component can be obtained directly from Equation (17):

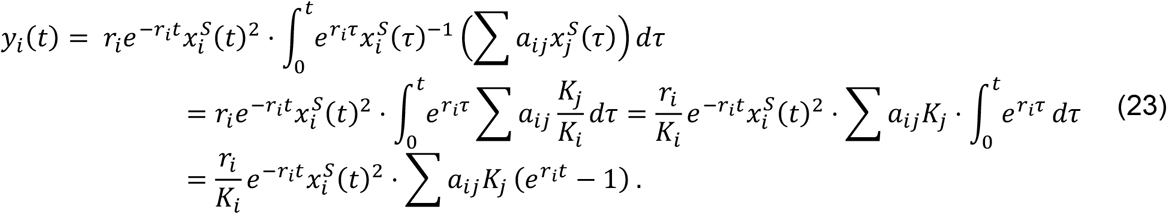

By substituting this expression into Equation (21), we obtain:

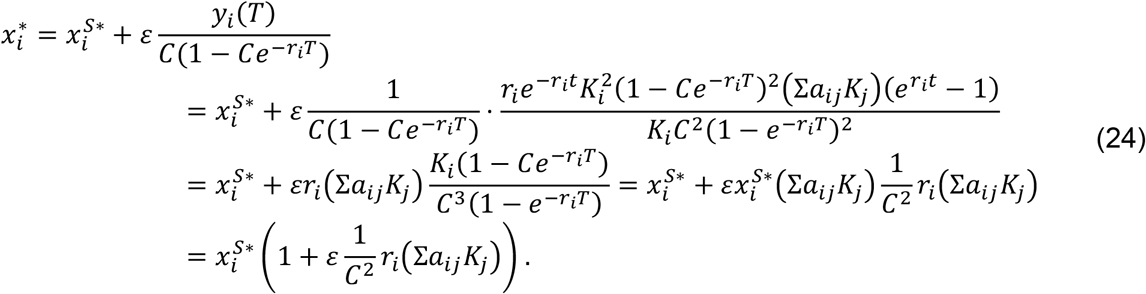

## 3. Results

### 3.1. Microbial abundance under recurrent disturbance depends primarily on *T*/log *C*, not on *T* and *C* separately

We begin by visualizing how individual species respond to a regularly recurrent disturbance. In **Figure 1**, we pick a representative community to demonstrate. For each species in the community, we plot its abundance profile on the (*T*, log *C*) plane, when grown either in isolation or in community.

**Figure 1.**
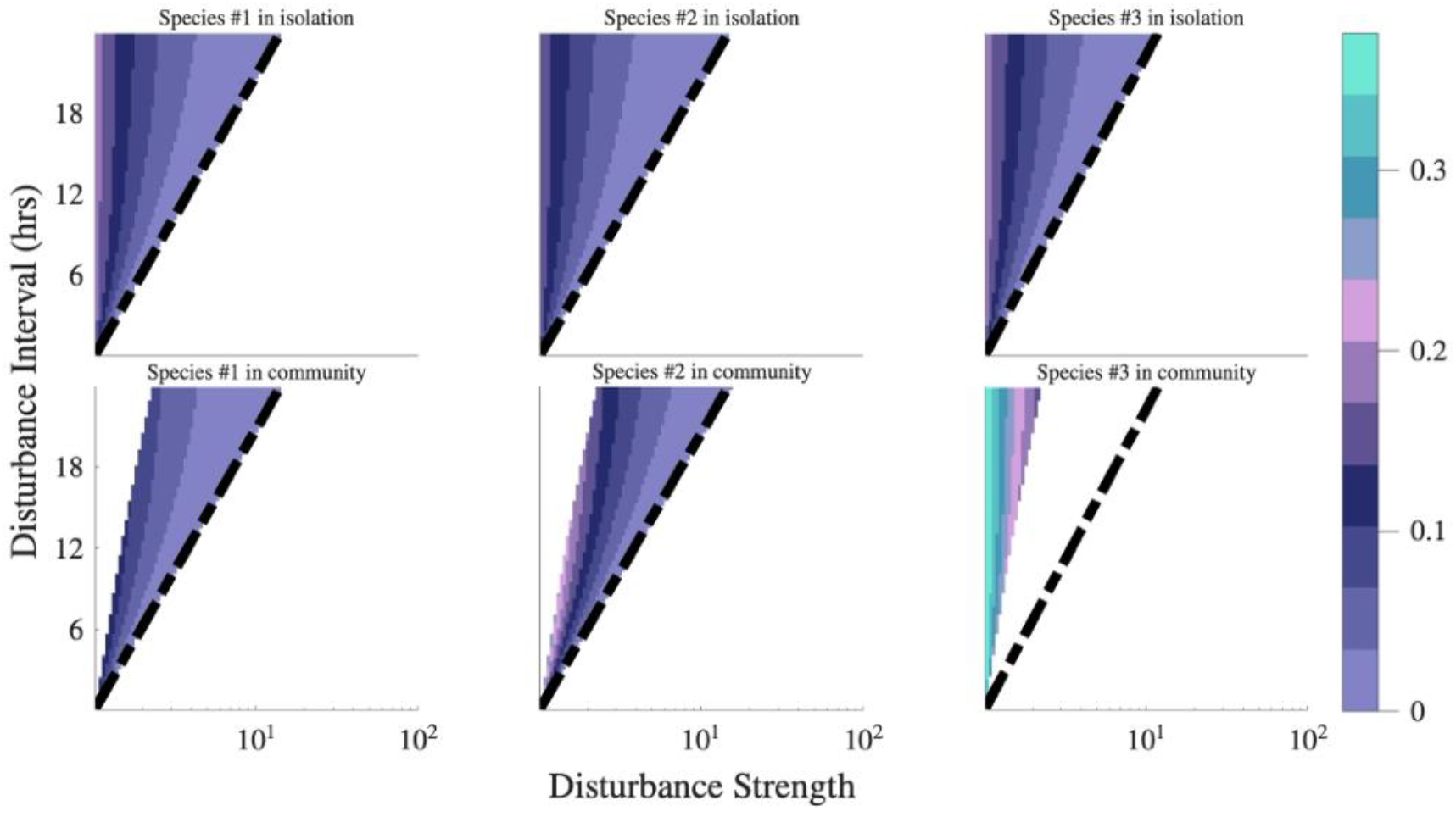
Microbial species abundance depends on the composite parameter *T*/log *C* in both isolation and community settings. Top row shows the abundance of three individual microbial species grown in isolation. Bottom row shows the abundance of the same species in a multi-species community. Columns correspond to different species. In each panel, color intensity indicates species abundance after 100 generations of *in-silico* growth under recurrent disturbance defined by dilution interval *T* (y-axis) and dilution factor *C* (x-axis, logscale). Experiments performed *in-silico* (see Methods). The region permitting growth exhibits linear boundaries in *T*/log *C* space. At low *T* and *C* close to continuous disturbance regime, abundance shows linear dependence on the composite parameter *T*/log *C*. At high *T* and *C*, abundance becomes vertically striped, depending primarily on *C*.

Several general patterns emerge. First, species exhibit an upper disturbance threshold: beyond sufficiently high *C*, they are unable to persist. This is consistent with expectation, since excessive disturbance intensity overwhelms any basal growth capacity. Interestingly, for every fixed *T*, the maximum survivable C values fall along a straight line in the (*T*, log *C*) plane, shown as a black dotted line in **Figure 1**.

Second, at very low disturbance intensities (i.e., low *C*), many species survive across the entire range of *T*; however, some species fail to persist when *C* becomes sufficiently small. In these cases, extinction arises not from the disturbance itself but from competitive exclusion: A competitor outcompetes the species and ultimately drives it to extinction. Again, for any fixed *T*, the minimum survivable *C* values align along a second straight line in the (*T*, log *C*) plane.

Third, the contour curves, that is, the level sets connecting all points in the abundance profile with the same final abundance, exhibit a strikingly simple structure. At low values of both *T* and *C*, these contour lines are approximately linear in the (*T*, log *C*) plane. However, as *T* increases, the contours begin to flatten, indicating a saturation regime in which increasing *T* further does not increase the abundance. This reveals a clear shift in the parameter that controls abundance: at low *T*, abundance varies primarily with the composite parameter *T*/log *C*, whereas at sufficiently high *T*, abundance depends predominantly on log *C* alone.

These observations suggests that species responses depend not on *T* and *C* independently, but primarily on their ratio, captured by *T*/log *C*. We confirmed this by showing that disturbance conditions with identical *T*/log *C* values exhibit dramatically lower variance in species abundance than the full dataset. This reduction in variability indicates that *T*/log *C* captures most of the impact of disturbance on species abundance.

### 3.2. Community abundance is independent of disturbance under weak interspecies interactions and similar growth rates

We analyze a simplified model amenable to theoretical treatment. In this model we assume weak interspecies interactions and similar growth rates across species (i.e., the assumptions of Neutral theory (Matthews & Whittaker, 2014)). Under these assumptions (detailed in Methods, Eq. 24), the end-of-cycle abundance of each species in the community is strictly proportional to its abundance when grown in isolation, with the proportionality factor depending only on species-specific parameters and independent of the disturbance parameters *T*.

Starting with equation (10) in the Methods section, we can calculate the stable abundance in isolation as

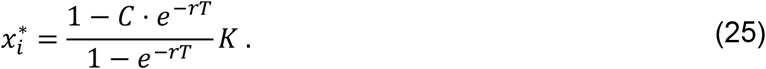

Note that although individual disturbances impact all species equally, the cumulative effects differ depending on species’ growth rate. For abundance to remain positive, the numerator of this fraction must be non-zero, establishing the viability condition *C*^−1^ − *e*^−*rT*^ > 0. Thus, *r* is a viable growth rate only if 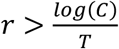. We define this critical growth rate as the *survival threshold* of each species. The survival threshold shows up as a line in the top panel of **Figure 1**. In the case of growth in a community, our theoretical calculation shows that abundance in a community is proportional to abundance in isolation, as shown in equation (24):

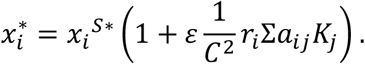

Here, 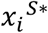 is the end of cycle abundance of a microbe grown in isolation, {*a*_*ij*_} represents the interspecies interaction strength and *K*_*i*_ is the carrying capacity of species *i*. This case reveals that the proportion is independent of disturbance frequency. We will show that a similar independence to disturbance is present in the general case.

### 3.3. Context-dependent interaction is largely insensitive to disturbance intensity or frequency

We ask whether interactions in a community are insensitive to disturbance when relaxing the assumptions of weak interspecies interactions and identical growth rates. We first define the context-dependent interaction (*I*_*CD*_) for a species *i* as:

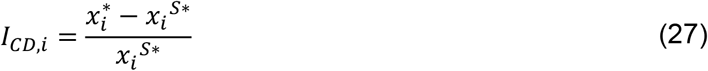

*I*_*CD*_ quantifies the net effect of the entire community context on species *i*. Unlike classical Lotka–Volterra interspecies interaction strengths (which are assumed constant between pairs), *I*_*CD*_ captures the net effect of the entire community context on species *i* and is therefore not necessarily constant. Importantly, *I*_*CD*_ is the observable property often measured in empirical investigations to quantify the overall impact of community interactions.

To test sensitivity of *I*_*CD*_ to disturbance parameters, we simulate 600 microbial communities across a wide range of disturbance regimes (*T* from 0.1–24 h, 120 values; *C* from 10^−1^–10^2^, 100 log-spaced values). We quantify sensitivity as the normalized absolute change in *I*_*CD*_ when *T* or *C* change (holding the other fixed):

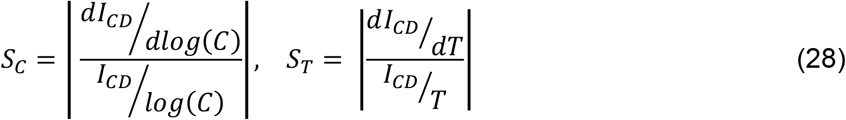

As shown in **Figure 2**, *I*_*CD*_ is largely insensitive to changes in disturbance frequency or intensity. In both *T*- and *C*-directions, most cases show normalized sensitivity <1. This demonstrates that context dependent interactions are generally robust to variation in disturbance regimes, although a subset of cases exhibits high sensitivity (as discussed below).

**Figure 2.**
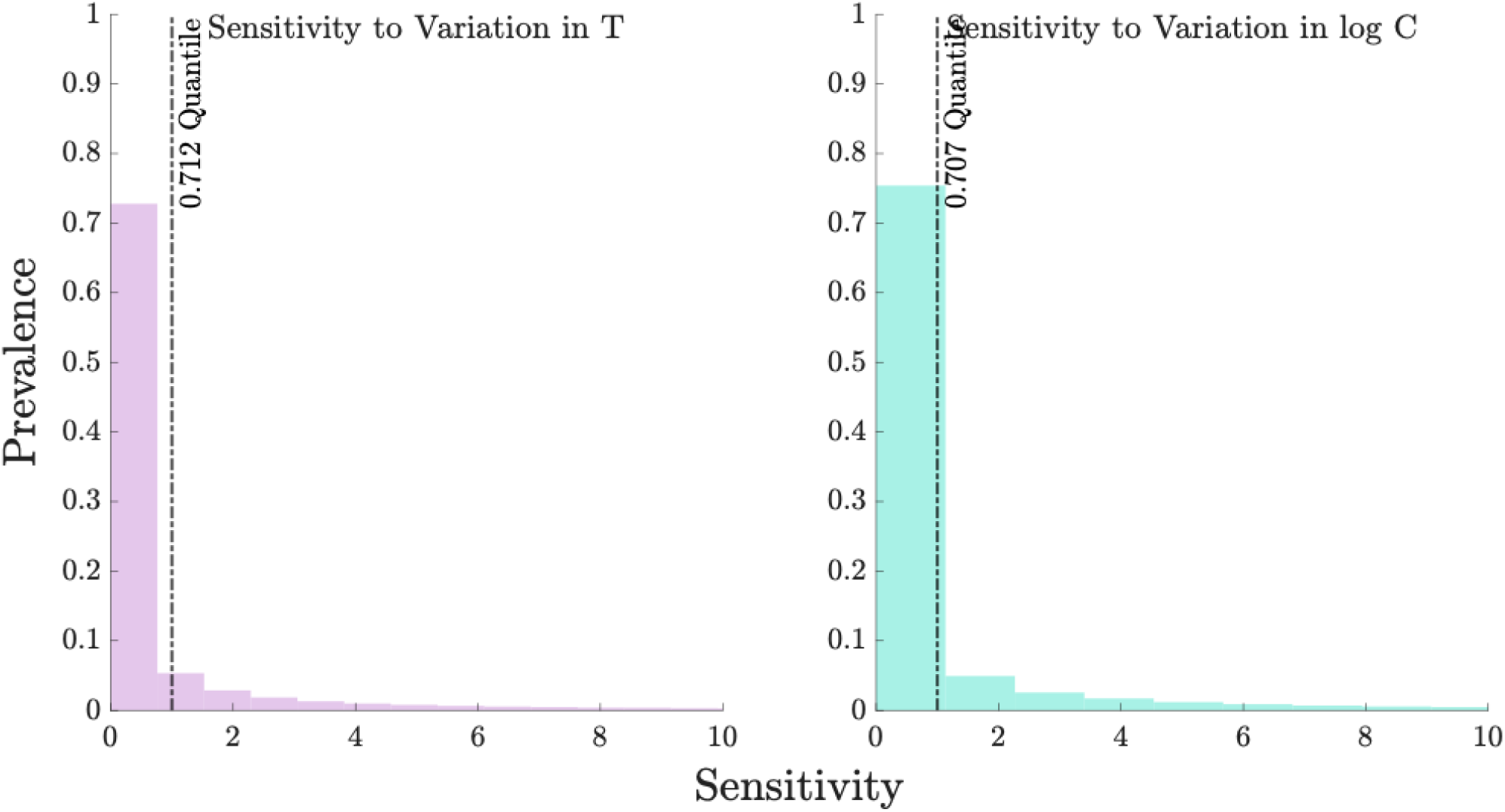
Context-dependent interactions (*I*_*CD*_) are largely insensitive to specific levels of disturbance parameters *T* and *C*. Normalized sensitivity of *I*_*CD*_ to incremental changes in *T* (left) and in *C* (right). Normalized sensitivity is defined as the absolute change in *I*_*CD*_ divided by the *I*_*CD*_ magnitude, analogous to a normalized derivative (See methods). Bar plots show the distribution across all tested conditions and across all communities. In both directions, >70% of cases exhibit low sensitivity (<1), indicating robustness to changes in disturbance regime, while a substantial fraction shows high sensitivity (>1) and will be examined in subsequent figures.

### 3.4. Context-dependent interactions sensitivity is confined to the transition region

We ask under what conditions *I*_*CD*_ become sensitive to disturbance. Inspecting equation (28) identifies two scenarios where elevated sensitivity is expected: (i) when *I*_*CD*_ approaches zero (no-interaction region), and (ii) when a species’ abundance approaches zero (survival threshold). We define the union of these two sets of boundaries as the transition region.

From **Figure 1**, species survival and abundance contours display linear patterns in *T*/log *C* space at low disturbance. Consequently, each (*T*, log *C*) condition can be represented by the ray originating from (0,0) on which it lies. Transition boundaries are also rays from the origin. We therefore quantify the location of any condition by its normalized angular distance to the nearest transition ray: the smallest angular separation between the two rays is normalized to the full angular span of conditions where the species can survive.

Using 600 simulated communities, we identified the top 5% most sensitive *I*_*CD*_ cases and measured their normalized angular distance to the transition region. As shown in **Figure 3**, nearly all highly sensitive cases occur in the vicinity of transition regions, demonstrating that elevated sensitivity is tightly confined to these boundaries.

**Figure 3.**
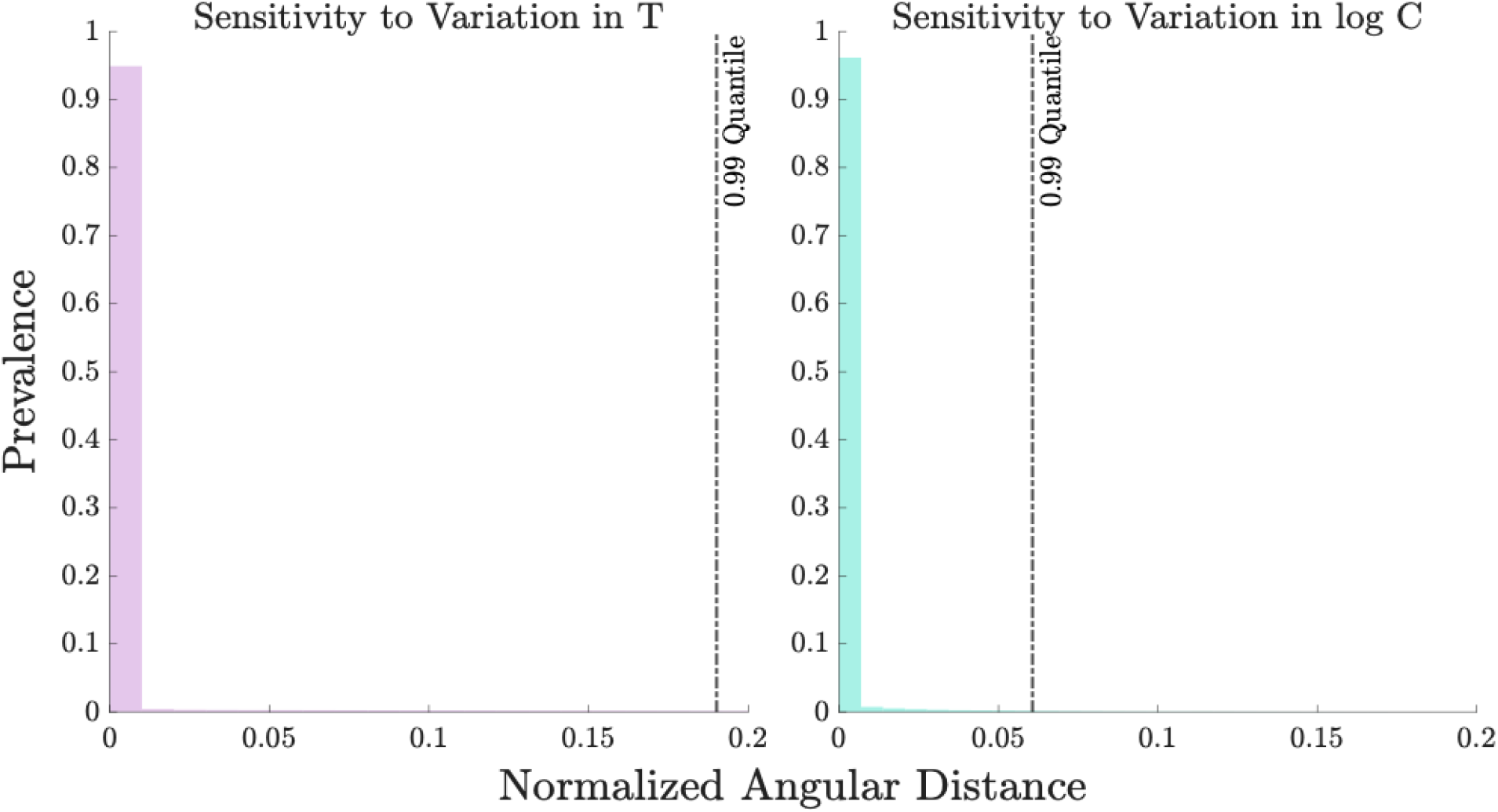
Sensitive context-dependent interactions (*I*_*CD*_) occur exclusively near the transition region. Distribution of normalized angular distance to the transition region for *I*_*CD*_ cases sensitive to changes in *T* (left) and to changes in *C* (right). The x-axis shows normalized angular distance to the transition region (defined in main text). In both panels, more than 95% of sensitive cases lie in the vicinity transition regions, confirming that sensitivity is confined to the transition region.

### 3.5. Irregularity in disturbance has negligible effect on stable community composition

Building on the disturbance-insensitivity of *I*_*CD*_ under regular regimes, we ask whether this insensitivity extends to irregular disturbances. To test this, we perform simulations in which irregularity is introduced in one parameter while keeping the other regular: either stochastic variation in disturbance interval *T* (irregular *T*, regular *C*) or in disturbance intensity *C* (irregular *C*, regular *T*), with variance of up to 50% of the baseline value.

As shown in **Figure 4**, community compositions under irregular disturbances closely match those under the corresponding regular regimes. In both cases, the vast majority of communities reach stable abundances with low Bray-Curtis dissimilarity (BC < 0.1) to the regular case. This demonstrates that final species abundances, and therefore *I*_*CD*_, are largely insensitive to the introduction of irregularity in either disturbance parameter. Note that here irregularity is applied to only one parameter at a time; however, we expect that these results extend to a fully irregular regime as well.

**Figure 4.**
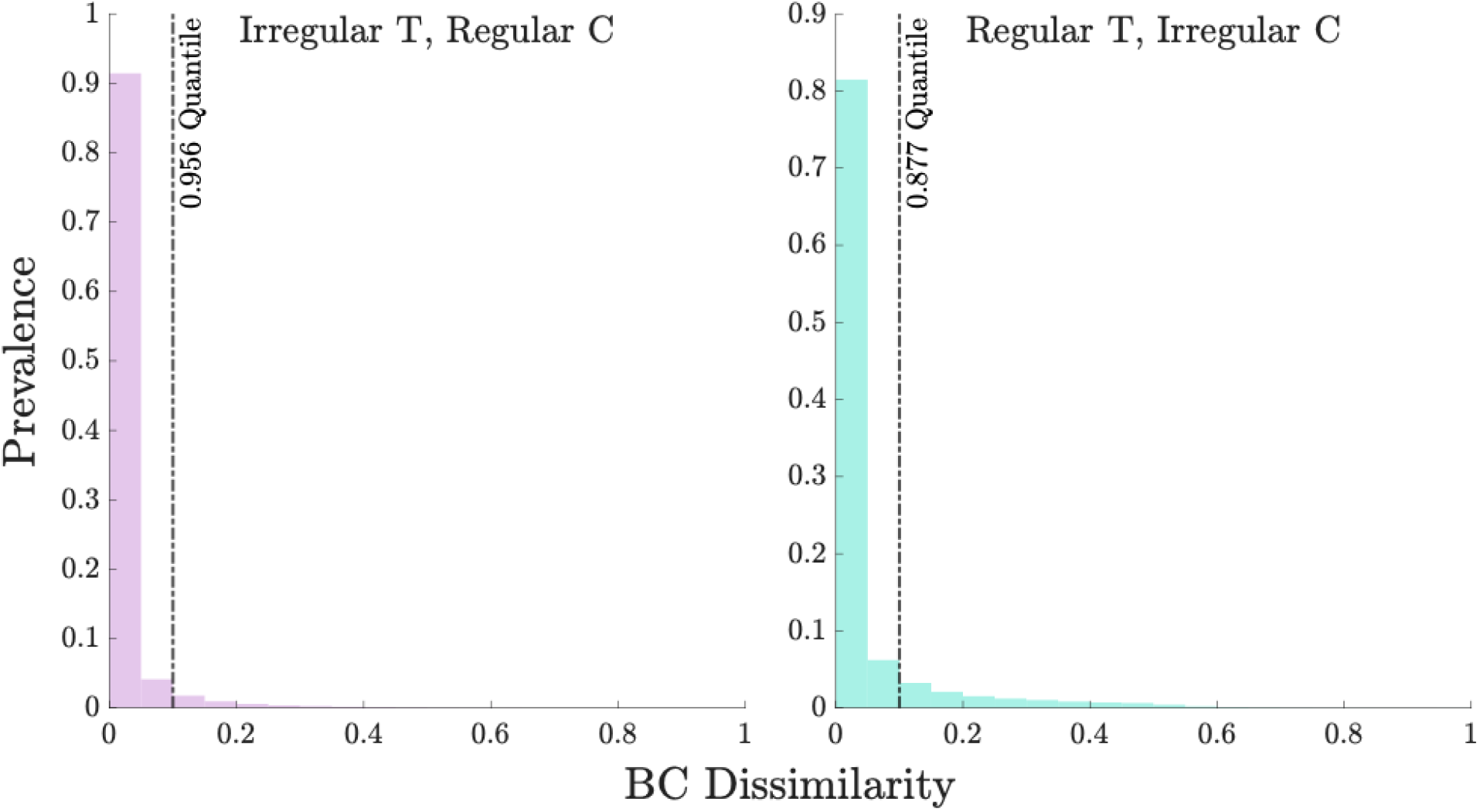
Community composition under irregular disturbances closely matches that under regular disturbances. Bray-Curtis dissimilarity (BC) between communities grown under (regular *C*, irregular *T*) versus regular *C* and *T* (left), under (irregular *C*, regular *T*) versus regular *C* and *T* (right). X-axis: Bray-Curtis dissimilarity of final stable abundances. Y-axis: Fraction of simulated communities at each dissimilarity value. In both panels, the large majority of communities show very low BC dissimilarity, indicating that species abundances stabilize at nearly identical levels under irregular and regular disturbance regimes.

The few communities that show substantial dissimilarity (BC > 0.1) are almost exclusively located near the transition region (**Figure S2**), consistent with the sensitivity patterns observed under regular disturbances. **Figure S1** further confirms that this close match between regular and irregular regimes holds across all tested levels of irregularity (10–50%), with only modest increases in outliers at the highest variance levels.

### 3.6. Uneven disturbance has negligible effect on stable community composition

We next ask whether community composition remains robust when disturbance intensity varies across species (uneven disturbance). Since unevenness in the disturbance interval *T* is not biologically well-defined in our system, we focus exclusively on introducing variation in the disturbance factor *C*. We hypothesize that uneven *C* would have minimal impact on stable community composition except near the transition region. To test this, we perform simulations in which each species experiences a random deviation (±10–50%) around the baseline *C* value while keeping *T* identical, and compare the resulting stable abundances to the even-disturbance case.

As shown in **Figure 5**, community compositions under even and uneven C regimes are highly similar. The large majority of communities exhibit low Bray-Curtis dissimilarity (BC < 0.1), indicating that uneven disturbance has negligible effect on final species abundances. **Figure S3** confirms that this robustness holds across all tested levels of unevenness (10–50%). As in previous analyses, the few communities showing substantial dissimilarity (BC > 0.1) are predominantly located near the transition region (**Figure S4**). This result extends the disturbance-insensitivity of community composition and *I*_*CD*_ to biologically relevant uneven disturbance scenarios.

**Figure 5.**
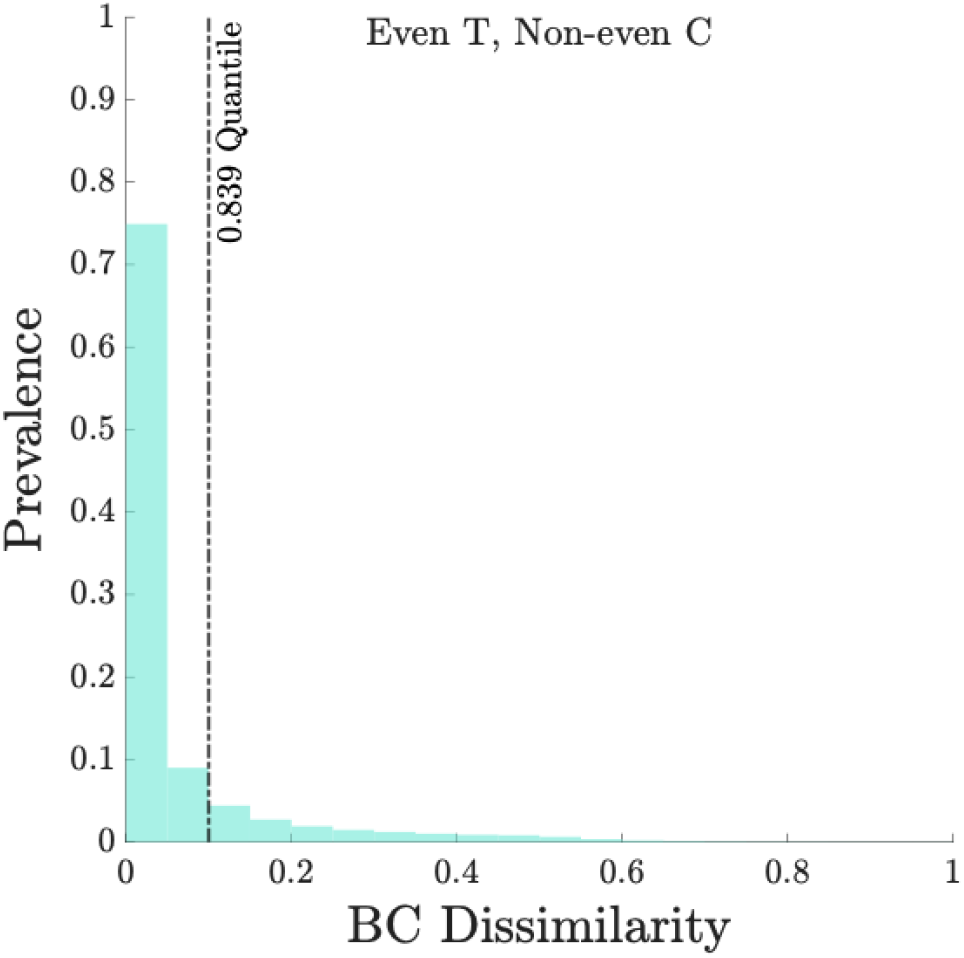
Community composition under uneven disturbances closely matches that under regular disturbances. Bray-Curtis dissimilarity (BC) between communities grown under even versus uneven dilution (*C*), where uneven disturbances introduce 10–50% variation in dilution factor across species. The large majority (>83%) of communities exhibit very low BC dissimilarity (<.01), demonstrating that species abundances under uneven *C* closely match those under even disturbance regimes. The x-axis is BC dissimilarity between even disturbance and un-even disturbance regimes for the same species. The y-axis shows the fraction of simulated communities at each level of dissimilarity.

### 3.7. Variation in community composition throughout the growth phase is minimal

To test whether our results hold not only at the end of the cycle but at any point during the growth phase, we compare Bray-Curtis dissimilarity between end-of-cycle abundances and abundances sampled at time points throughout the inter-disturbance growth phase across all stable communities and disturbance regimes.

As shown in **Figure 6**, community composition is highly stable throughout the growth phase. More than 99% of cases exhibit very low Bray-Curtis dissimilarity (BC < 0.1) between end-of-cycle and mid-cycle (or any intermediate) abundances. This demonstrates that the disturbance-independent patterns we observed are not specific to the end-of-cycle snapshot but hold continuously during growth. The rare outliers with elevated dissimilarity (BC > 0.1) occur almost exclusively near the transition region (**Figure S5**), consistent with previous findings.

**Figure 6.**
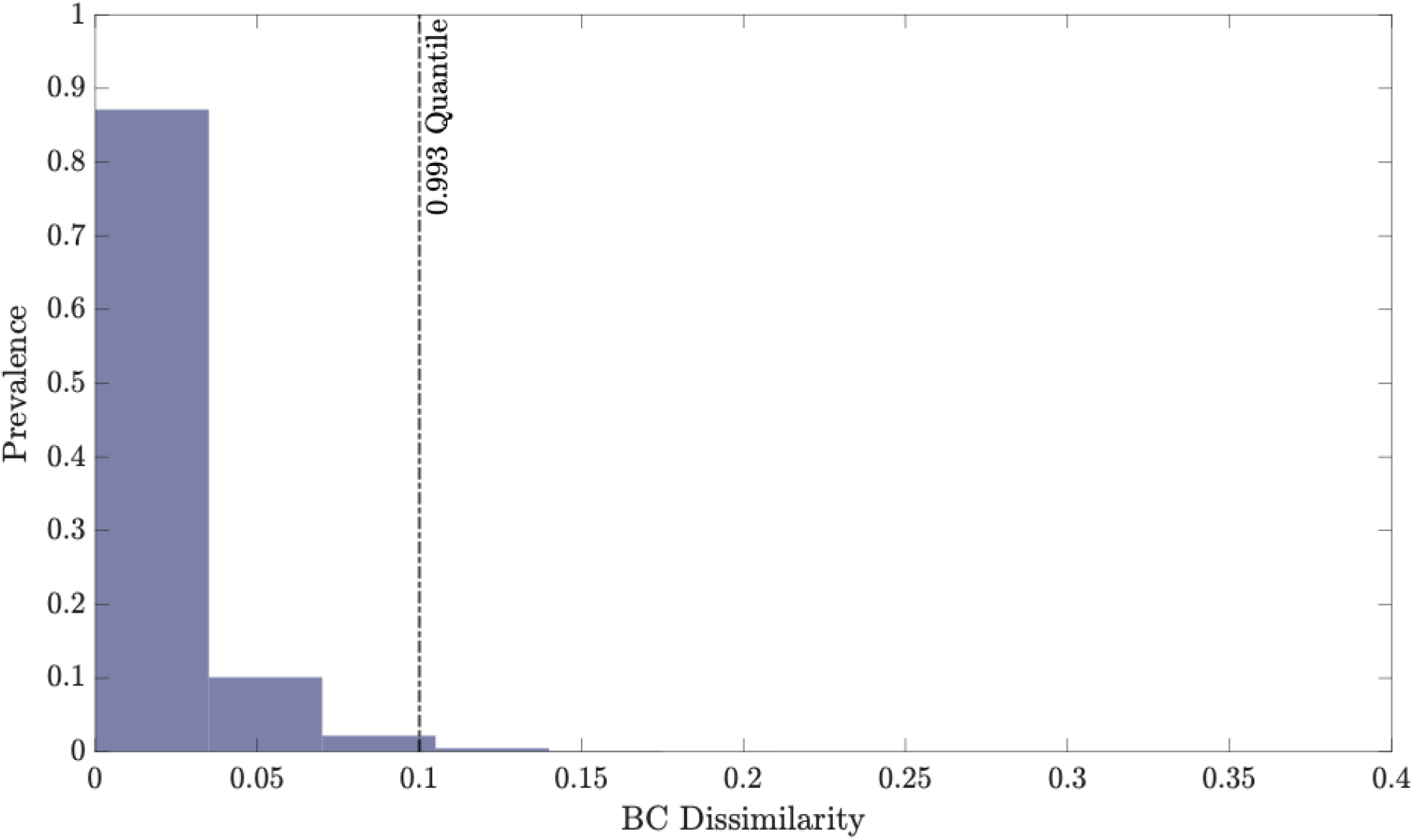
Microbial composition changes only slightly through the growth phase. Distribution of Bray–Curtis dissimilarity (BC) between end-of-cycle abundances and abundances at random time points during the growth phase, across all communities and disturbance regimes. More than 99% of cases show BC < 0.01, indicating that species abundances remain nearly constant relative to each other throughout the cycle.

## 4. Discussion

Environmental disturbances are ubiquitous in microbial ecosystems and are widely assumed to strongly shape community composition and microbial interactions within communities. Here, we examined how recurrent disturbances influence these properties. Contrary to common intuition, we find that increasing disturbance intensity or frequency has minimal impact on community composition and context-dependent interactions (*I*_*CD*_), except in regimes strong enough to drive species extinctions. We first demonstrated this robustness under regular, even disturbances using end-of-cycle abundances, then extended the results to irregular and uneven disturbance regimes, and finally showed that community composition remains stable throughout the entire inter-disturbance growth phase. Together, these findings reveal a striking robustness of microbial communities to recurrent disturbances.

A consistent pattern emerging from both our analytical derivations and numerical simulations is that the effects of disturbance intensity (*C*) and frequency (1/*T*) are captured by a single composite parameter *T*/log *C*, rather than by the two parameters independently. This reduces the two-dimensional disturbance space to a one-dimensional axis. Species survival depends only on whether its intrinsic growth rate exceeds a critical threshold set by this composite parameter. Likewise, species abundances and context-dependent interactions show minimal variation among regimes that share the same value of *T*/log *C*. Motivated by this reduction, we quantified differences between disturbance regimes using a normalized angular distance in (*T*, log *C*) space (based on the arctangent of the ray from the origin), enabling direct comparison of community responses across widely varying conditions.

Our conclusions are based on a deliberately simplified mathematical model chosen for its analytical tractability and ability to systematically explore disturbance effects. We began with weak interactions and identical growth rates, then relaxed these assumptions in large-scale simulations. Sensitivity analyses show that the key qualitative patterns, disturbance robustness and confinement of sensitivity to the boundary region, hold across a wide range of species richness (2–10 species; **Figures S6** and **S7**), growth rates, carrying capacities, and interaction strengths. This robustness to model details supports the generality of the observed principles within the framework studied.

Future work could extend this framework to consumer–resource models or other formulations that relax the fixed-interaction assumption of the Lotka–Volterra model. To preserve analytical clarity, such extensions should be introduced incrementally, beginning with minimal species-resource networks and simple consumption topologies. Ultimately, these theoretical predictions can be tested experimentally in controlled microbial communities to validate the observed resilience and scaling relationships.

Our results also contextualize the classical focus in microbial ecology on disturbance magnitude. While sufficiently strong disturbances can drive extinctions and thereby alter diversity and composition, we show that within sub-extinction regimes, community composition and context-dependent interactions remain highly robust. Extinction (or proximity to extinction thresholds) is the primary pathway through which recurrent disturbances reshape microbial communities.

Collectively, these findings reframe disturbance as a weaker force than commonly assumed. Rather than viewing microbial communities as fragile systems easily restructured by environmental fluctuations, our results highlight robustness as an intrinsic property of multispecies systems. Experimentally, this implies that minor procedural variations, such as differences in dilution intervals, incubation times, or sampling delays, typically have negligible effects on community composition, as long as extinction thresholds are not crossed. More broadly, this resilience suggests that preserving species persistence, rather than eliminating environmental variability, may be the key to maintaining stable microbial communities in both natural and engineered ecosystems.

## Acknowledgments

This work was supported by the National Science Foundation (NSF MCB) under Grant No. 2430384 and by a SI-GECS grant from the Schiller Institute for Integrated Science and Society at Boston College.

## Code availability

All the relevant codes are available on GitHub at https://github.com/vahramian/env_disturbance.

## Supplementary Information

**Figure S1.**
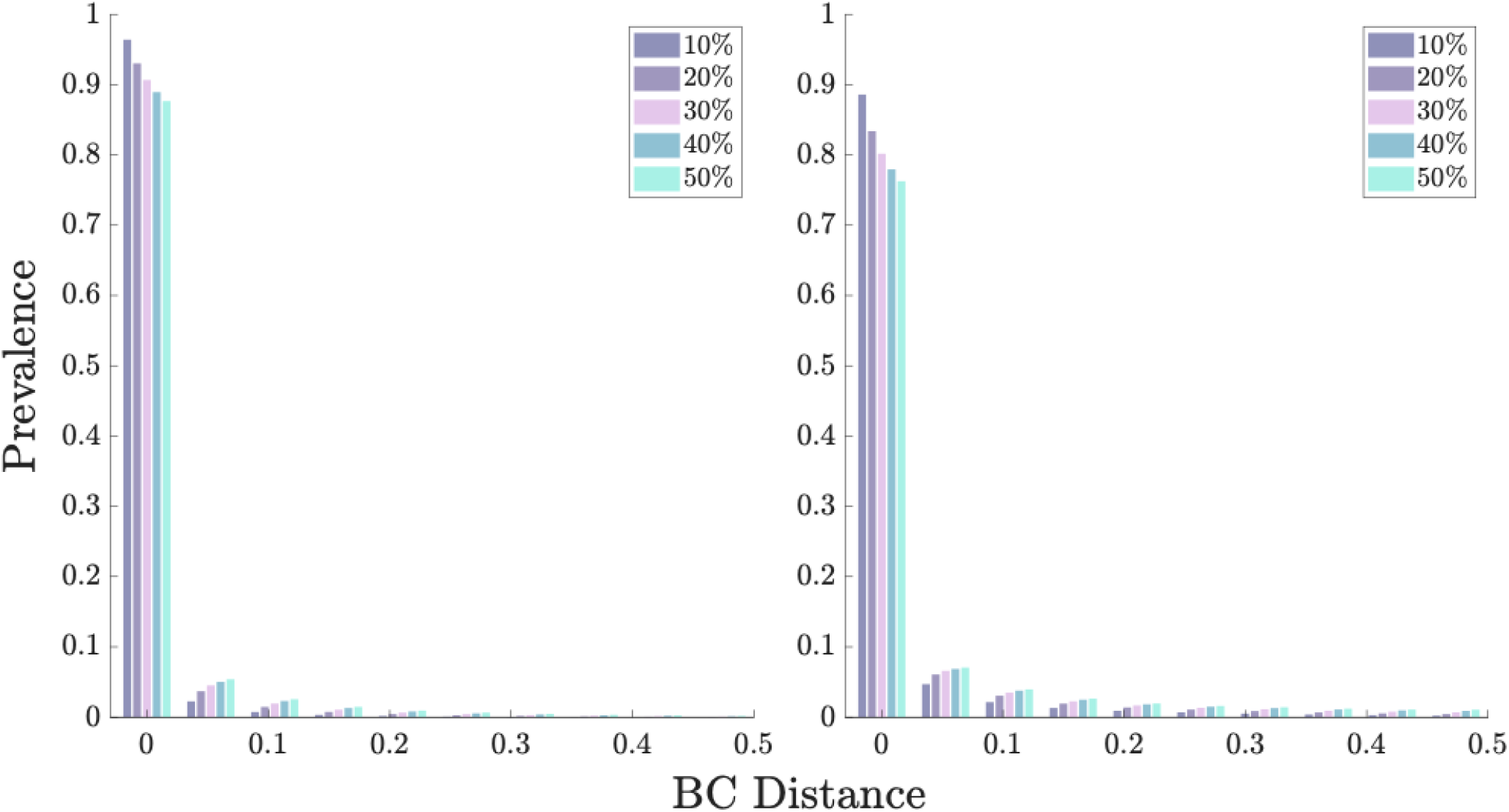
Effect of increasing irregularity on community composition. Vertical bars represent the five irregularity levels. Regardless of irregularity level (10–50%), the vast majority of communities exhibit very low BC dissimilarity, confirming that the close match between regular and irregular disturbance regimes is robust to the degree of irregularity.

**Figure S2.**
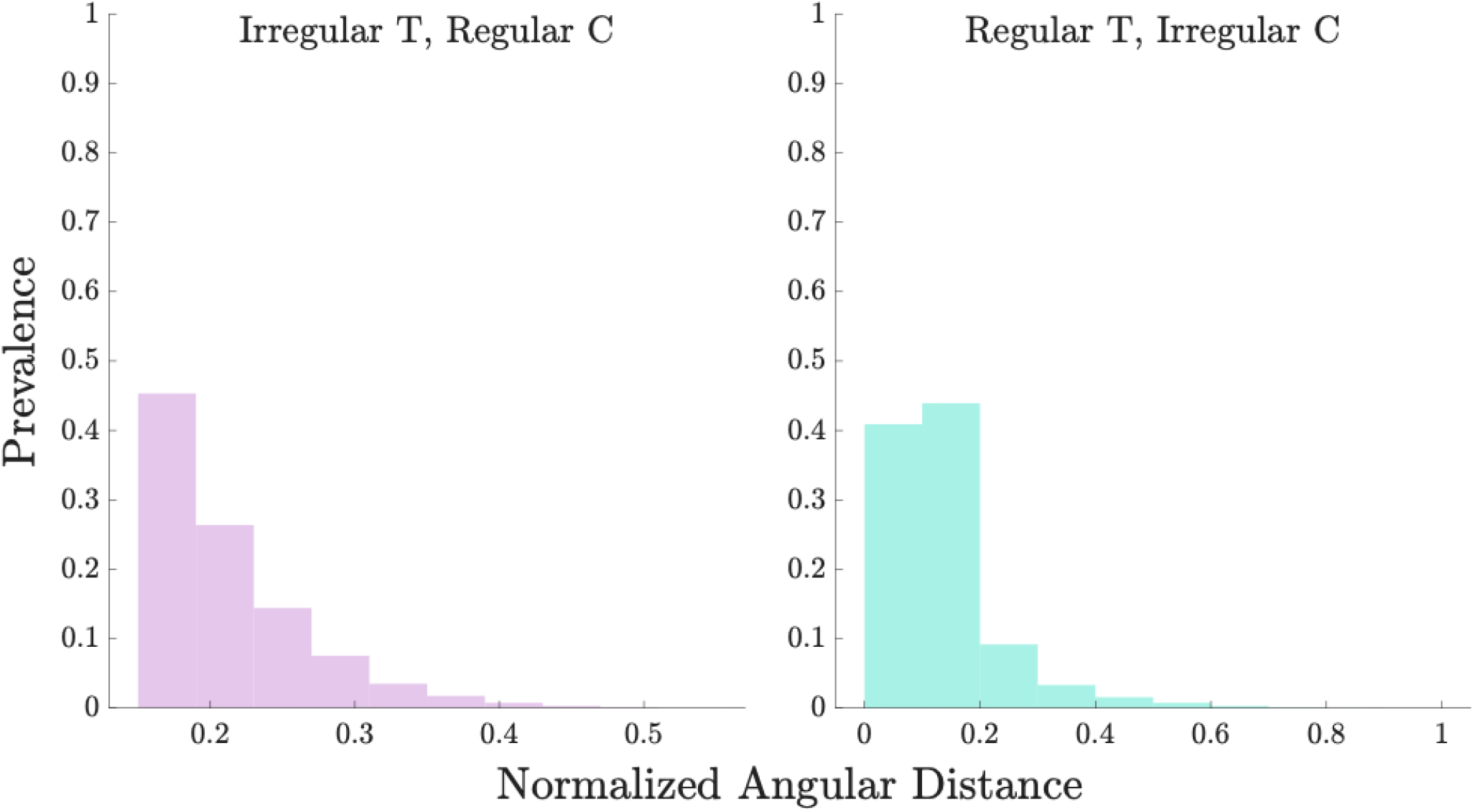
Cases with significant dissimilarity under irregular disturbances occur near the transition region. Normalized angular distance to the boundary region for communities with Bray-Curtis dissimilarity (BC) > 0.1 when *T* is irregular (left), or when *C* is irregular (right). The x-axis is the normalized angular distance to the nearest transition region (as defined in **Figure 3**).

**Figure S3.**
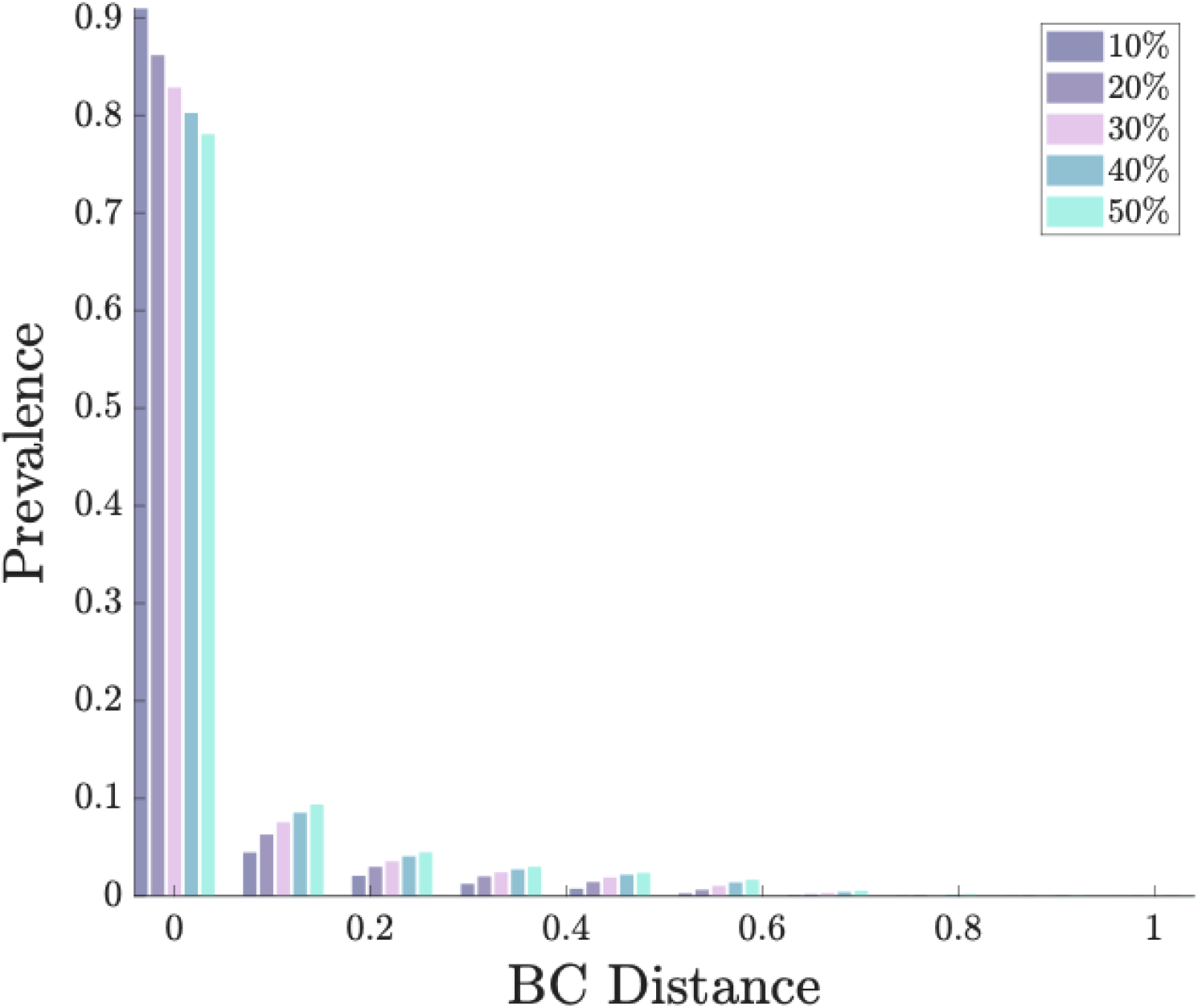
Effect of increasing unevenness on community composition. Bray-Curtis dissimilarity (BC) between communities grown under even versus uneven dilution (*C*), stratified by degree of unevenness (10%, 20%, 30%, 40%, 50% variation across species). Regardless of the degree of unevenness (10–50%), the large majority of communities show very low BC dissimilarity, confirming that community composition remains close to the even-disturbance case at all tested levels of uneven *C*. The x-axis is the BC dissimilarity of final stable abundances between even and uneven cases. They-axis is the fraction of simulated communities at each level of dissimilarity.

**Figure S4.**
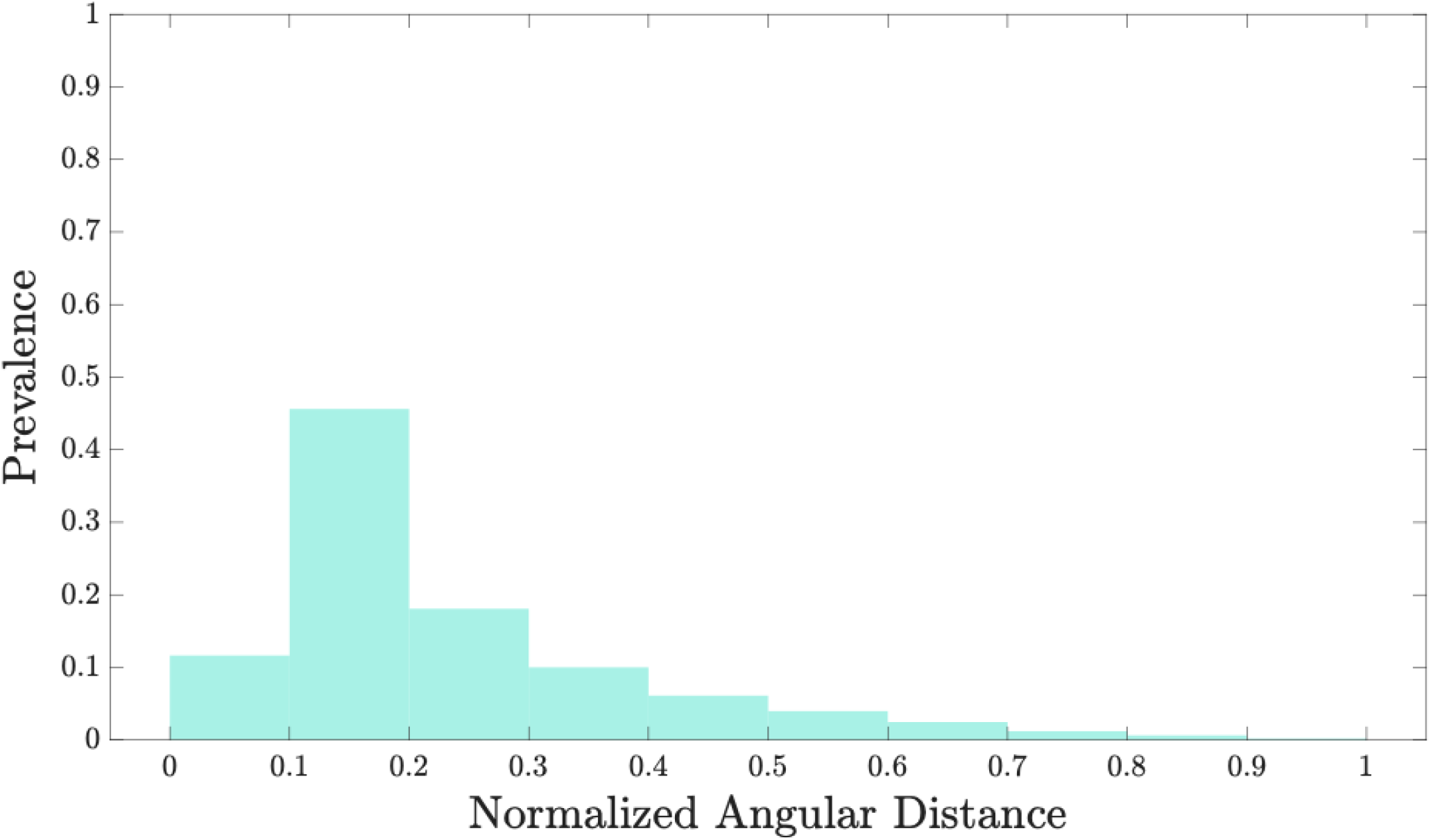
High-dissimilarity cases under uneven disturbances are confined to the transition region. Normalized angular distance to the boundary region for communities showing Bray-Curtis dissimilarity (BC) > 0.1 between even and uneven *C* regimes. The x-axis is the normalized angular distance to the nearest transition region (as defined in **Figure 3**).

**Figure S5.**
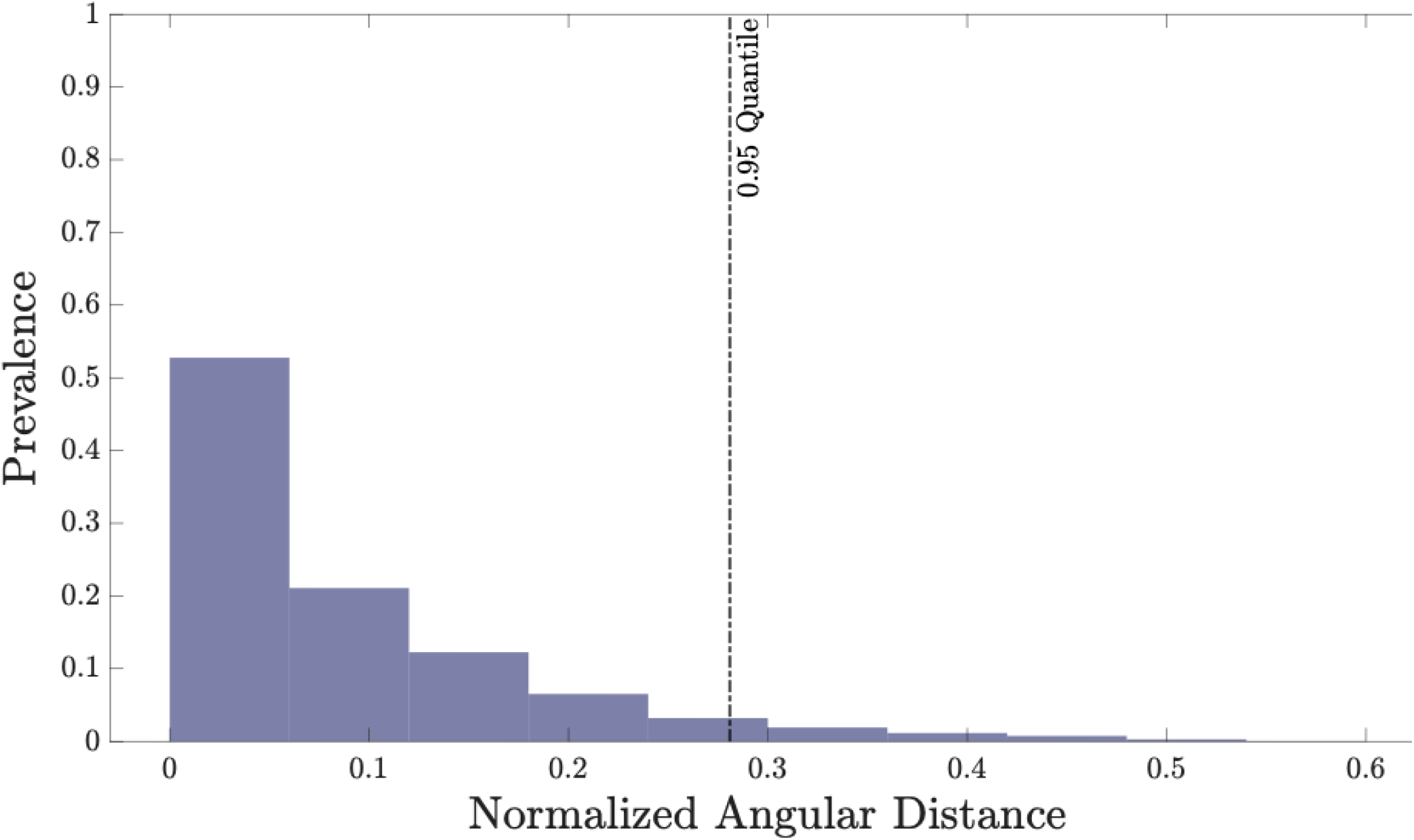
High-dissimilarity cases during the growth phase occur near the boundary region. Normalized angular distance to the boundary region for communities showing elevated Bray-Curtis dissimilarity (BC > 0.1) between end-of-cycle and intermediate growth-phase abundances.

**Figure S6.**
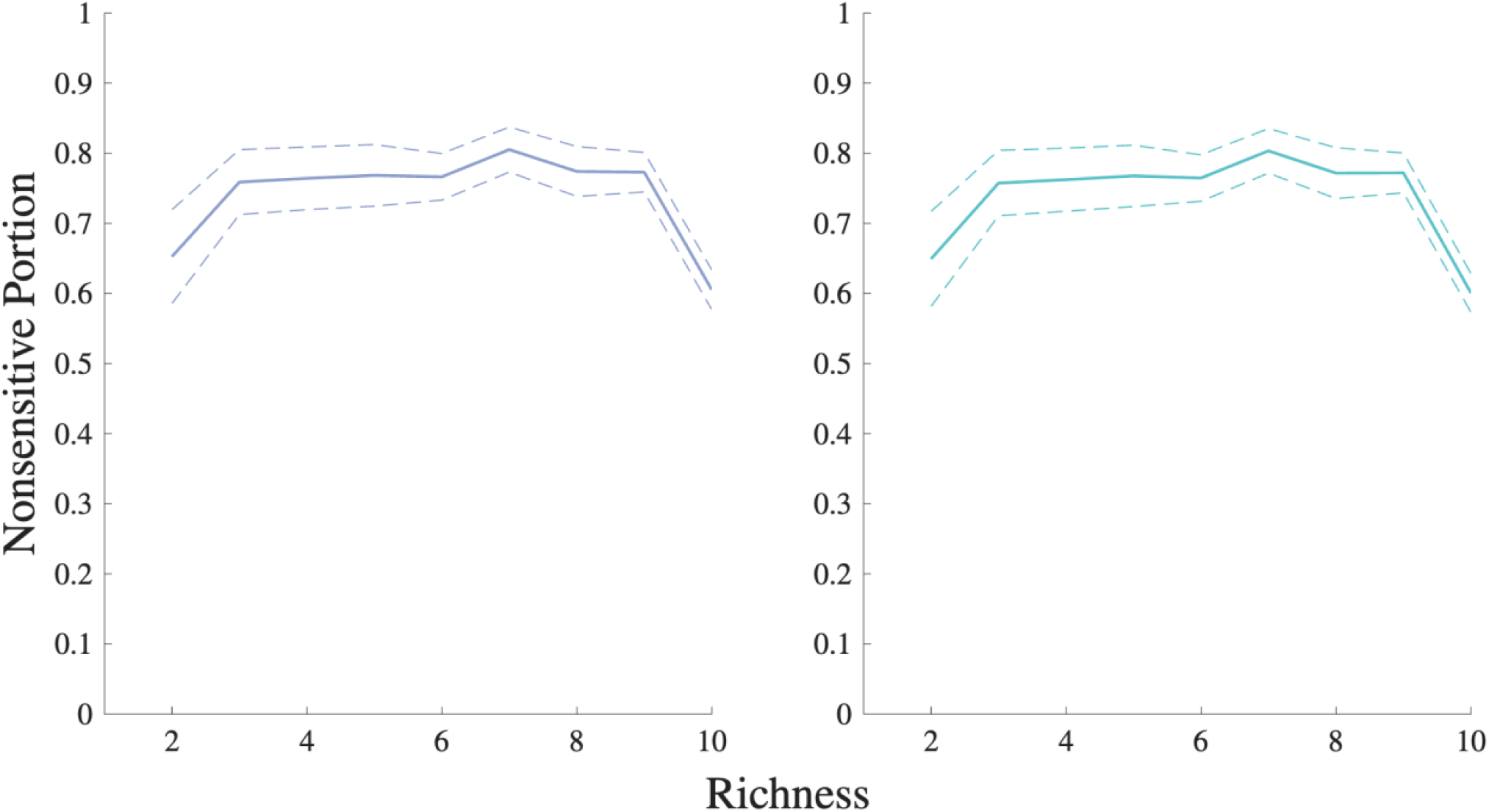
Proportion of disturbance-insensitive *I*_*CD*_ cases is independent of species richness. Proportion of cases with normalized sensitivity <1 to changes in *T* (left) and in *C* (right). Lines show mean values across simulations, and dashed lines are confidence intervals; richness ranges from 2 to 10 species. The y-axis is the proportion of cases with low sensitivity at each level of richness.

**Figure S7.**
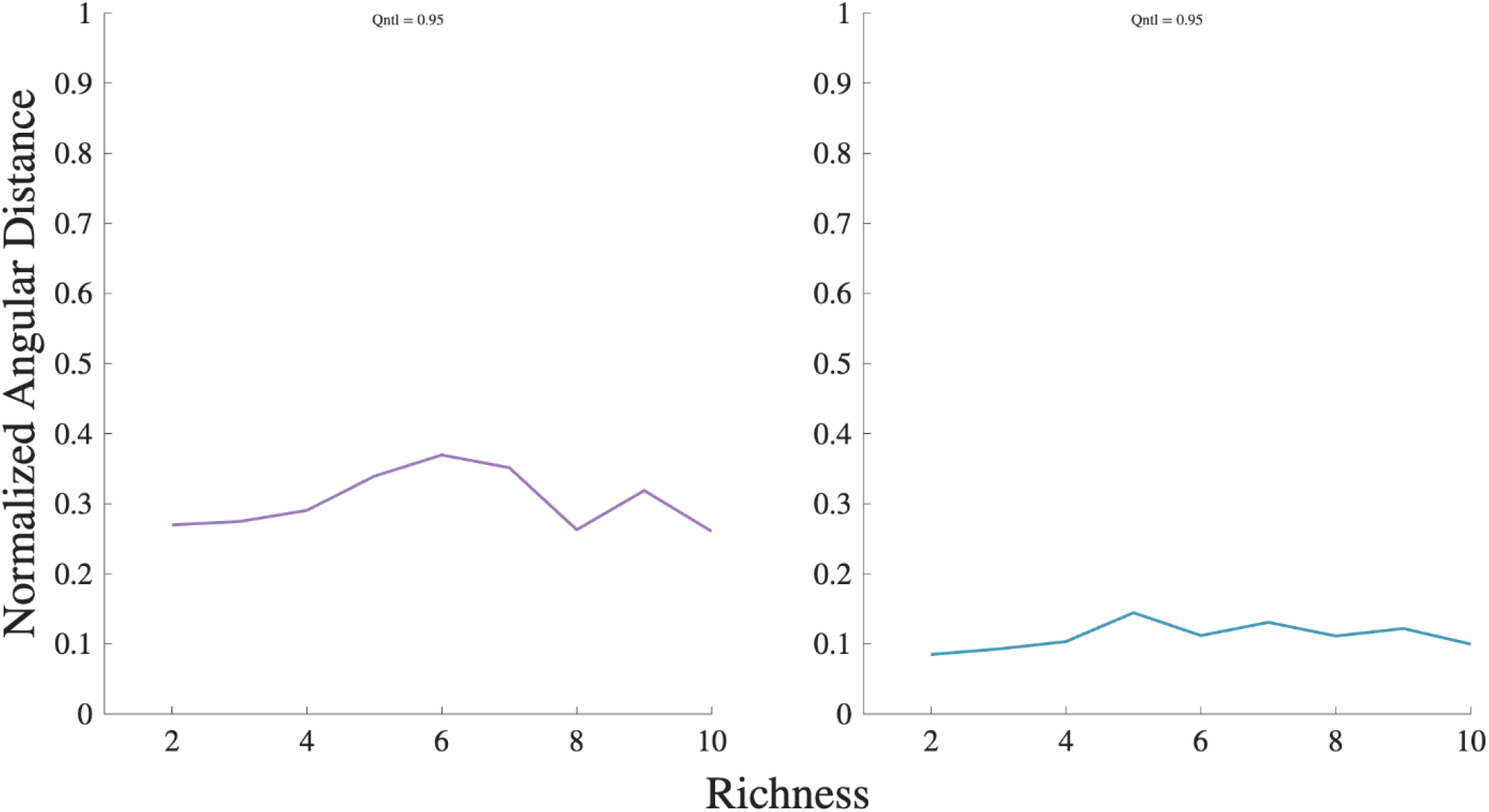
Sensitive *I*_*CD*_ cases remain close to the transition region regardless of species richness. Normalized angular distance to the boundary region for *I*_*CD*_ cases sensitive to changes in *T* (left) or in *C* (right). The x-axis shows species richness (2–10). The y-axis is the normalized angular distance to the nearest transition region. Lines represent the distance of the top 5% outliers across simulations at each richness level. The consistently low distances confirm that sensitivity is driven by proximity to the transition region rather than community richness.

## References

Banitz, T., Chatzinotas, A., & Worrich, A. (2020). Prospects for Integrating Disturbances, Biodiversity and Ecosystem Functioning Using Microbial Systems. Frontiers in Ecology and Evolution, 8. 10.3389/fevo.2020.00021

Berga, M., Székely, A. J., & Langenheder, S. (2012). Effects of Disturbance Intensity and Frequency on Bacterial Community Composition and Function. PLoS ONE, 7(5), e36959. 10.1371/journal.pone.0036959

Buckley, H. L., Day, N. J., Lear, G., & Case, B. S. (2021). Changes in the analysis of temporal community dynamics data: a 29-year literature review. PeerJ, 9, e11250. 10.7717/peerj.11250

Buma, B. (2021). Disturbance ecology and the problem of n = 1: A proposed framework for unifying disturbance ecology studies to address theory across multiple ecological systems. Methods in Ecology and Evolution, 12(12), 2276–2286. 10.1111/2041-210X.13702

Burman, E., & Bengtsson-Palme, J. (2021). Microbial Community Interactions Are Sensitive to Small Changes in Temperature. Frontiers in Microbiology, 12. 10.3389/fmicb.2021.672910

Dal Bello, M., Lee, H., Goyal, A., & Gore, J. (2021). Resource–diversity relationships in bacterial communities reflect the network structure of microbial metabolism. Nature Ecology & Evolution, 5(10), 1424–1434. 10.1038/s41559-021-01535-8

Dedrick, S., Akbari, M. J., Dyckman, S. K., Zhao, N., Liu, Y.-Y., & Momeni, B. (2021). Impact of Temporal pH Fluctuations on the Coexistence of Nasal Bacteria in an in silico Community. Frontiers in Microbiology, 12. 10.3389/fmicb.2021.613109

Fujikawa, H., & Sakha, M. Z. (2014). Prediction of Competitive Microbial Growth in Mixed Culture at Dynamic Temperature Patterns. Biocontrol Science, 19(3), 121–127. 10.4265/bio.19.121

Gallardo-Navarro, O., Aguilar-Salinas, B., Rocha, J., & Olmedo-Álvarez, G. (2024). Higher-order interactions and emergent properties of microbial communities: The power of synthetic ecology. Heliyon, 10(14), e33896. 10.1016/j.heliyon.2024.e33896

Gonze, D., Coyte, K. Z., Lahti, L., & Faust, K. (2018). Microbial communities as dynamical systems. Current Opinion in Microbiology, 44, 41–49. 10.1016/j.mib.2018.07.004

Hu, J., Barbier, M., Bunin, G., & Gore, J. (2025). Collective dynamical regimes predict invasion success and impacts in microbial communities. Nature Ecology & Evolution, 9(3), 406–416. 10.1038/s41559-024-02618-y

Kurkjian, H. M., Akbari, M. J., & Momeni, B. (2021). The impact of interactions on invasion and colonization resistance in microbial communities. PLOS Computational Biology, 17(1), e1008643. 10.1371/journal.pcbi.1008643

Lee, H., Bloxham, B., & Gore, J. (2023). Resource competition can explain simplicity in microbial community assembly. Proceedings of the National Academy of Sciences, 120(35). 10.1073/pnas.2212113120

Mancuso, C. P., Lee, H., Abreu, C. I., Gore, J., & Khalil, A. S. (2021). Environmental fluctuations reshape an unexpected diversity-disturbance relationship in a microbial community. ELife, 10. 10.7554/eLife.67175

Matthews, T. J., & Whittaker, R. J. (2014). Neutral theory and the species abundance distribution: recent developments and prospects for unifying niche and neutral perspectives. Ecology and Evolution, 4(11), 2263–2277. 10.1002/ece3.1092

Momeni, B., Xie, L., & Shou, W. (2017). Lotka-Volterra pairwise modeling fails to capture diverse pairwise microbial interactions. ELife, 6. 10.7554/eLife.25051

Nguyen, J., Lara-Gutiérrez, J., & Stocker, R. (2021). Environmental fluctuations and their effects on microbial communities, populations and individuals. FEMS Microbiology Reviews, 45(4). 10.1093/femsre/fuaa068

Philippot, L., Griffiths, B. S., & Langenheder, S. (2021). Microbial Community Resilience across Ecosystems and Multiple Disturbances. Microbiology and Molecular Biology Reviews, 85(2). 10.1128/MMBR.00026-20

Schimel, J. P. (2018). Life in Dry Soils: Effects of Drought on Soil Microbial Communities and Processes. Annual Review of Ecology, Evolution, and Systematics, 49(1), 409–432. 10.1146/annurev-ecolsys-110617-062614

Shade, A., Peter, H., Allison, S. D., Baho, D. L., Berga, M., Bürgmann, H., Huber, D. H., Langenheder, S., Lennon, J. T., Martiny, J. B. H., Matulich, K. L., Schmidt, T. M., & Handelsman, J. (2012). Fundamentals of Microbial Community Resistance and Resilience. Frontiers in Microbiology, 3. 10.3389/fmicb.2012.00417

Stein, R.R., Bucci, V., Toussaint, N.C., Buffie, C.G., Ratsch, G. Pamer, E.G., Sander, C., Xavier, J. (2013). Ecological Modeling from Time-Series Inference: Insight into Dynamics and Stability of Intestinal Microbiota. PLOS Computational Biology 9(12), e1003388. 10.1371/journal.pcbi.1003388.

Song, H.-S., Cannon, W., Beliaev, A., & Konopka, A. (2014). Mathematical Modeling of Microbial Community Dynamics: A Methodological Review. Processes, 2(4), 711–752. 10.3390/pr2040711

Venturelli, O. S., Carr, A. V, Fisher, G., Hsu, R. H., Lau, R., Bowen, B. P., Hromada, S., Northen, T., & Arkin, A. P. (2018). Deciphering microbial interactions in synthetic human gut microbiome communities. Molecular Systems Biology, 14(6). 10.15252/msb.20178157

